# Zika virus-induces metabolic alterations in fetal neuronal progenitors that could influence in neurodevelopment during early pregnancy

**DOI:** 10.1101/2022.08.24.505080

**Authors:** Javier Gilbert-Jaramillo, Ujang Purnama, Zoltán Molnár, William S. James

## Abstract

Neuronal progenitor subtypes have distinct fate restrictions regulated by time-dependent activation of energetic pathways. Thus, the hijacking of cellular metabolism by Zika virus (ZIKV) to support its replication may contribute to damage in the developing fetal brain. Here, we showed that ZIKV replicates differently in two glycolytically distinct hiPSC-derived neuronal progenitors that correspond to early and late progenitors in the forebrain. This differential replication alters the transcription of metabolic genes and upregulates the glycolytic capacity of progenitor subtypes. Analysis using Imagestream® revealed that, during early stages of infection, ZIKV replication in early progenitors increases lipid droplet abundance and decreases mitochondrial size and membrane potential. During later stages infection, early progenitors show increased subcellular distribution of lipid droplets, whilst late progenitors show decreased mitochondria size. The finding that there are hi-NPC subtype-specific alterations of cellular metabolism during ZIKV infection may help to explain the differences in brain damage over each trimester.

## Introduction

Intrauterine development during the first trimester of pregnancy includes a series of orchestrated cellular processes, mainly cell division and differentiation, which give origin to primitive tissue and organs(1). The brain develops from a neural plate that forms a tube. The inner lining of this tube contains the germinal zone where most of the progenitor cells are situated(2). Neuronal progenitor cells at different stages of proliferation and differentiation exhibit specific activation and shifts between the main cellular metabolic pathways (glycolysis, glucose and fatty acid oxidation)(3–5). The shift from cytosolic glucose metabolism to mitochondrial glucose oxidation and the activation of fatty acid oxidation are thought to signal for differentiation of quiescent neuronal progenitors to proliferative neuronal progenitors(3–5).

Maternal malnutrition during pregnancy may alter the nutrient supply and metabolism of the fetal brain imposing severe consequences to normal development(1,6). These consequences are comparable and potentially contribute to those observed during infections with neurotropic viruses such as Zika virus (ZIKV)(7). The maternal-fetal circulation allows ZIKV to reach the developing fetal brain where its infection and metabolic hijacking of neuronal cells potentially underpin the anatomical and physiological damage observed in new-borns. This damage is mainly observed after infection during the first trimester(8,9) and is not observed to the same extent when maternal infection occurs during mid and late trimesters(10,11), after the accelerated fetal brain expansion has occurred.

There are no studies investigating whether ZIKV infection imposes differential metabolic stress in neuronal progenitor subtypes as a potential cause for the specific brain damage over each trimester. Thus, we investigated the metabolic stress imposed by ZIKV infection in human induced pluripotent stem cell (hiPSCs) derived cortical neuronal progenitors (hi-NPCs) that resembled the two main subtypes of early progenitors of the forebrain. Here, we show that these progenitor subtypes differ in their rate of glucose and fatty acid metabolism and that this is differentially exploited by ZIKV. ZIKV infection increases the glycolytic capacity of progenitor subtypes and, to a larger extend, of those who metabolise less glucose. Notably, few changes in the intracellular distribution and size of lipid droplets and the mitochondrial homeostasis are observed in early progenitors at different stages of ZIKV replication.

## Results

### Differentiation of human cortical neuronal progenitors rescues metabolically distinct neuronal progenitor subtypes

Differentiation of cortical neuronal progenitors (hi-NPCs) using a 2D adaptation from Shi et al., 2012; Robbins et al., 2018(12,13) (Fig. 1A), produces, at different time-points, hi-NPCs with distinct morphology (Fig. 1B and 1C). Based on the time of differentiation, these cells were called early and late hi-NPCs. Early hi-NPCs were rounded and exhibited a large cytoplasm and a large nucleus with few membrane projections (Fig. 1B), whilst “star-shaped” late hi-NPCs displayed abundant projections with a small cytoplasm and nucleus (Fig. 1C). Morphological characterization by flow cytometry showed that differentiation of early and late hi-NPCs cultures produced subpopulations of brain cell types (neuronal precursors, glia, and neurons). These populations did not significantly differ (p ≥ 0.05) between hi-NPCs cultures (Fig. 1D). Screening of neuronal progenitors’ markers showed no significant differences (p ≥ 0.05) in the expression of most of the proteins of interest between hi-NPCs subtypes with Pax6 showing a significantly higher expression in late hi-NPCs (Fig. 1E). Positive staining of neuronal progenitors’ markers was validated by confocal imaging (Fig. 1F and Extended Data Fig. 1).

**Fig.1.**
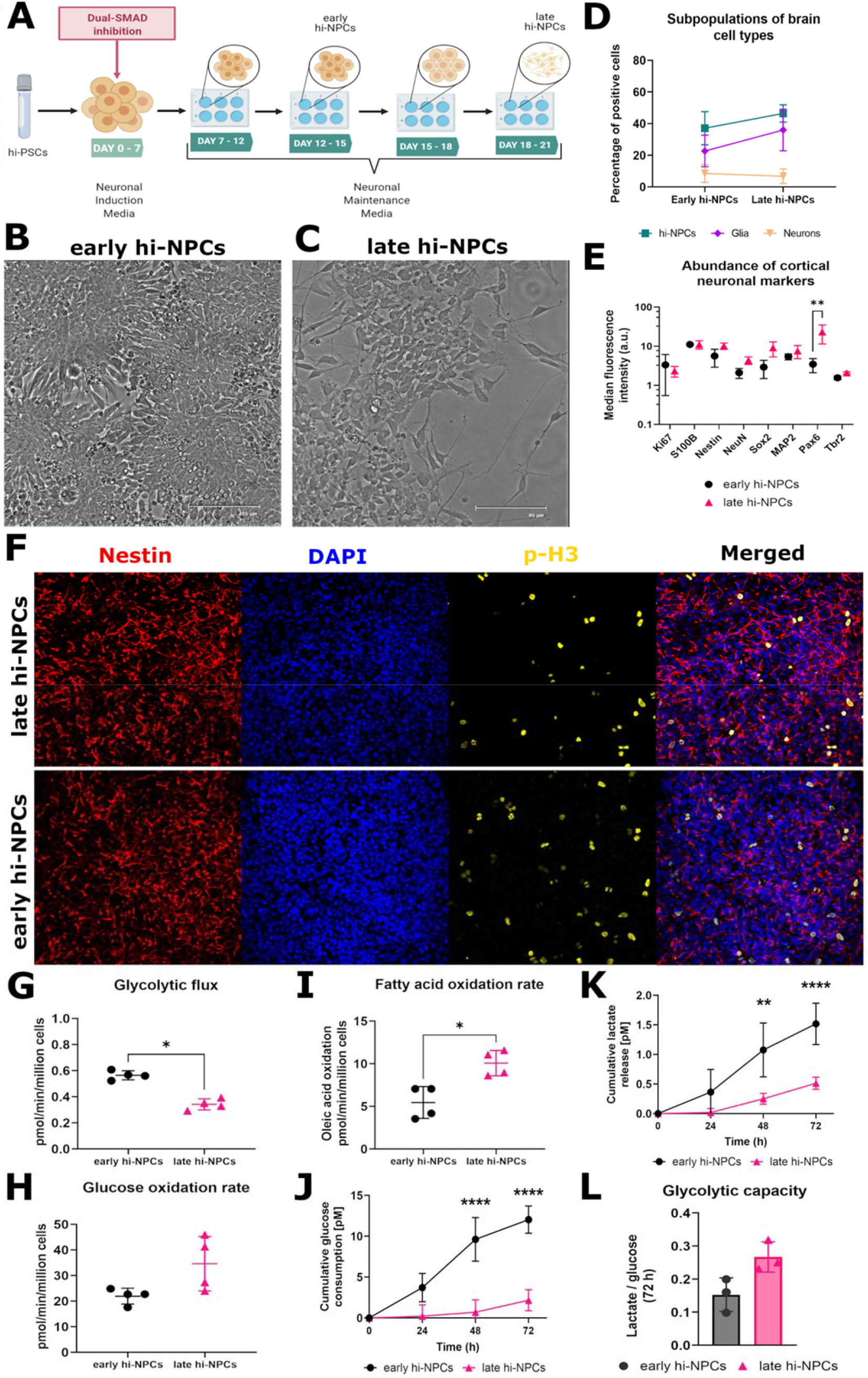
hiPSC cortical differentiation produces neuronal progenitor subtypes. (A) Schematic of 2D differentiation of hiPSCs to cortical neuronal progenitors (hi-NPCs). Brightfield images showing differences in morphology between early (B) and late hi-NPCs (C). Scale bars of 125 μm and 95 μm, respectively. (D) Dot plot showing the percentage summary of brain cell types in early and late hi-NPCs. n = 1 cortical differentiation done in triplicates for each of three independent donors’ cell lines. Significance was calculated by two-way ANOVA with Šidák’s multiple comparisons post hoc test. (E) Dot plot showing the relative median fluorescence intensity of a panel of conventional early cortical neuronal progenitor markers in early and late hi-NPCs. n = 1 cortical differentiation of two independent donors’ cell lines each conducted in four replicates. Significance was calculated by a mixed-effects model with Šidák correction. (F) Representative confocal images (10x) of the detection of markers of cellular proliferation phospho-Histone3 (p-H3) and of *in vitro* cortical neuronal progenitors nestin in early and late hi-NPCs. Scale bar: 100 μm. Intracellular radioactive tracing showing the metabolic remodelling of the (G) glycolytic flux, (H) glucose oxidation rate and, (I) fatty acid oxidation rate between early and late hi-NPCs. n = 2 cortical differentiations each measured at two different time-points, 10 and 14 h post plating, in five replicates for a donor cell line. Significance was calculated by non-parametric two-tailed Mann-Whitney U test. Dot plots showing the estimated cumulative glucose consumption (J) and lactate release (K) in hi-NPC subtypes cultured over 72 h. (L) Bar graphs displaying the glycolytic capacity of per hi-NPC subtype at 72 h post-culture. n = 1 cortical differentiation measured in triplicates for three independent donors’ cell lines. Significance was calculated by Two-way ANOVA with Šidák’s multiple comparisons post hoc test. Error bars display mean ± SD. Significance is shown when *p < 0.05, ** p < 0.01, *** p < 0.001, **** p < 0.0001.

Cellular metabolism is crucial during differentiation of neuronal progenitor cells(14). We traced intracellular short-term metabolism (between 10- and 14-hours post-passage) using radioactive-labelled substrates and found that the metabolic output from hi-NPC cultures differ from each other. Analysis of glucose processing showed a significantly lower glycolytic flux (p = 0.0286) in late compared to early hi-NPCs (Fig. 1G) and no differences in the mitochondrial glucose oxidation (Fig. 1H). The estimation of mitochondrial lipid oxidation by measuring the oxidation rate of oleic acid showed to be significantly higher in late compared to early hi-NPCs (p = 0.0286) (Fig. 1I). To assess potential changes in cellular metabolism induced by ZIKV replication we periodically examined the glucose consumption and lactate release. Early hi-NPCs, compared to late hi-NPCs, showed a significant increase in the cumulative levels of glucose consumption (p < 0.0001) and lactate release (p = 0.0037 and p = 0.0006, respectively) at 48 h and 72 h (Fig. 1J and 1K). However, the glycolytic capacity, estimated as a ratio of pico Molar [pM] of lactate released per [pM] of glucose consumed per cell, showed not to be significantly different between hi-NPC subtypes (Fig. 1L).

### Zika virus (ZIKV) differentially replicates in neuronal progenitor subtypes inducing subtype-specific alterations

ZIKV challenges in hi-NPC subtypes (Fig. 2A) may elucidate contributing mechanisms for the distinct brain damage induced by ZIKV infection during the first trimester. Thermal decay of ZIKV during the time of infection showed not to be significantly decreased (Fig. 2B). Both hi-NPC subtypes showed susceptibility to ZIKV infection with differences in the replication rates. RNA quantification showed that late hi-NPCs accumulate significantly more transcripts of ZIKV in a time-dependent manner than early hi-NPC. At 48 h.p.i., late hi-NPCs showed a 1.77-fold increase in intracellular copies of vRNA compared the early hi-NPCs. This further increased to a 1.98-fold change at 72 h.p.i. (Fig. 2C). Analysis of the intracellular accumulation of the non-structural NS1 and the virion-associated Envelope (Env) viral proteins by imaging flow cytometry revealed greater NS1 levels in late hi-NPCs at 56 h.p.i. (p = 0.0241) (Fig. 2D). However, we detected low percentages of NS1 positive cells. Detection of Env showed higher percentages of detection with no significant differences between hi-NPC subtypes (Fig. 2E). Infectious virus released by cells was significantly higher in late hi-NPCs exclusively at 56 h.p.i. compared to early hi-NPCs (late:early ratio of 2.70) (Fig. 2F). Cell viability following ZIKV challenge, assessed using the CCK8 tetrazolium assay, revealed significant cell death of both hi-NPC subtypes at later stages of ZIKV replication (p ≤ 0.0014) (Fig. 2G) yet, to different rates.

**Fig. 2.**
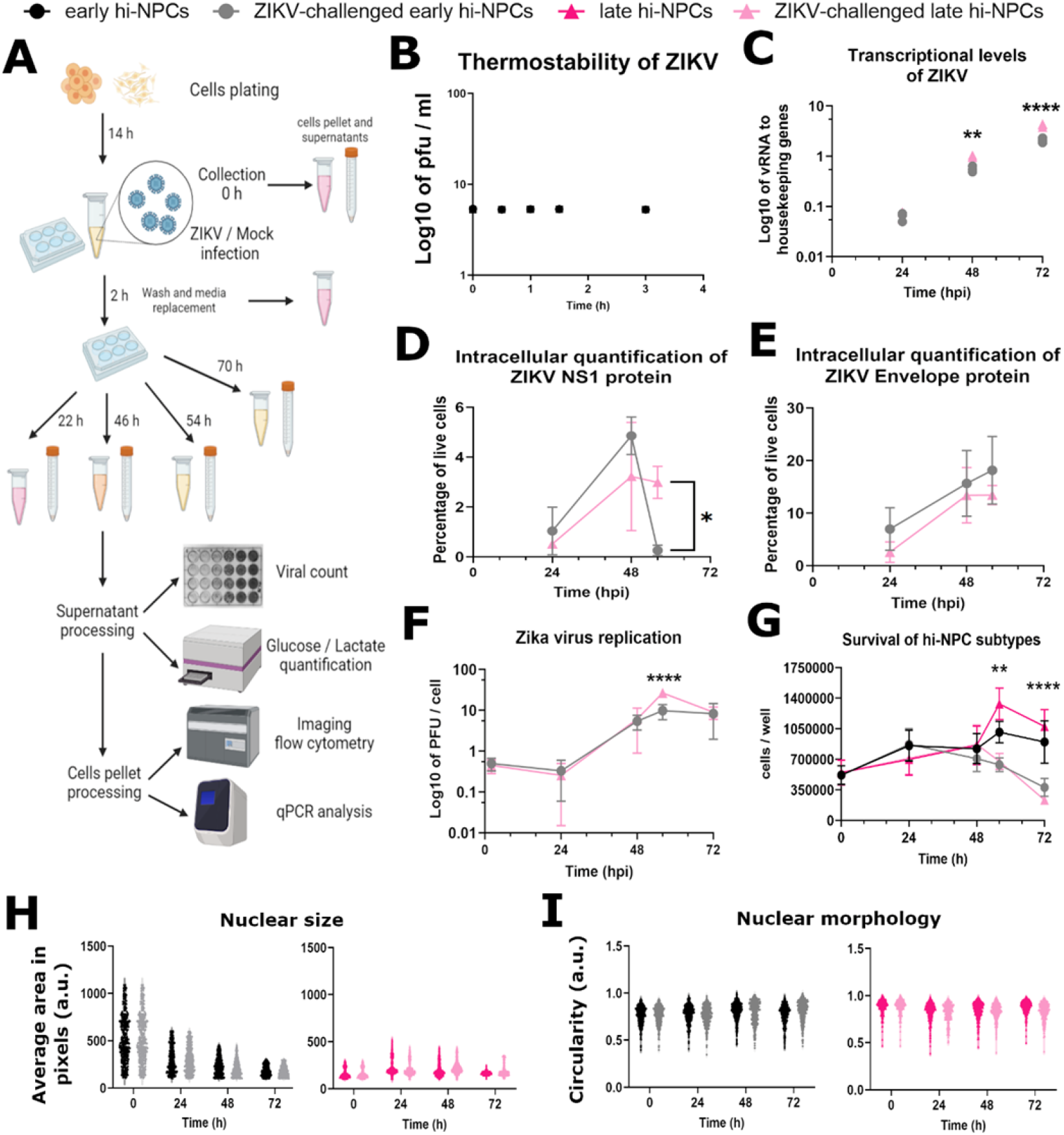
ZIKV exhibits differential intracellular assembly rates causing subtype-specific cellular damage in neuronal progenitors. (A) Schematic 2D representation of the protocol conducted to investigate the effects of hi-NPC subtypes exposure to ZIKV. (B) Dot plot showing the stability of ZIKV at 37 °C. n = 3 experiments titrated by plaque assay in triplicates. Significance was calculated by a mixed-effects model with Dunnet’s multiple comparisons test. (C) Transcriptional levels of ZIKV relative to housekeeping genes. qPCR conducted in four replicates. Dots displaying the values of each donor cell line. Percentage of live cells containing (D) ZIKV NS1 protein and (E) ZIKV Envelope protein. A minimum of 500 *in focus* cells were analyzed per donor cell line out of 10000 cells recorded. n = 1 viral challenge conducted in duplicates of three independent donors’ cell lines. Significance was calculated by Two-way ANOVA with Šidák’s multiple comparisons post-hoc test. (F) Dot plot showing extracellular release of ZIKV infectious particles by hi-NPCs. ZIKV quantification was done by plaque assay in triplicates. n = 2 viral challenges conducted in duplicates of three independent donors’ cell lines. Significance was calculated by mixed-effects model with Šidák correction. (G) Dot plots showing the cell viability of hi-NPCs exposed to ZIKV and mock controls. n = 1 viral challenge conducted in triplicate of one donor cell line. Significance was calculated by mixed-effects model with Holm-Šidák correction. Violin plots displaying the mean value for ≥ 150 nuclei per condition. Nuclei stained with DAPI showing (H) nuclear size and (I) nuclear morphology of hi-NPCs. Significance was calculated by mixed-effects model with Šidák’s multiple comparisons test. Error bars display mean ± SD. Significance is shown when *p < 0.05, ** p < 0.01, *** p < 0.001.

In addition, ZIKV challenges reflected changes in nuclear morphology that were distinct between neuronal progenitor subtypes (Extended Data Fig. 2 and Fig 3). However, these changes were potentially exclusive to infected cells thus, measurements of nuclear damage represented as size and morphology did not show significant differences between non-challenged and challenged conditions. At 24 h.p.i., ZIKV-challenged early hi-NPCs showed a reduction in nuclear size whilst the opposite effect was observed in ZIKV-challenged late hi-NPCs at 48 h.p.i. (Fig. 2H). Nuclear circularity, measured by roundness of the area stained by DAPI, was increased in ZIKV-challenged early hi-NPCs and decreased in ZIKV-challenged late hi-NPCs exclusively at 72 h.p.i. (Fig. 2I).

**Fig. 3.**
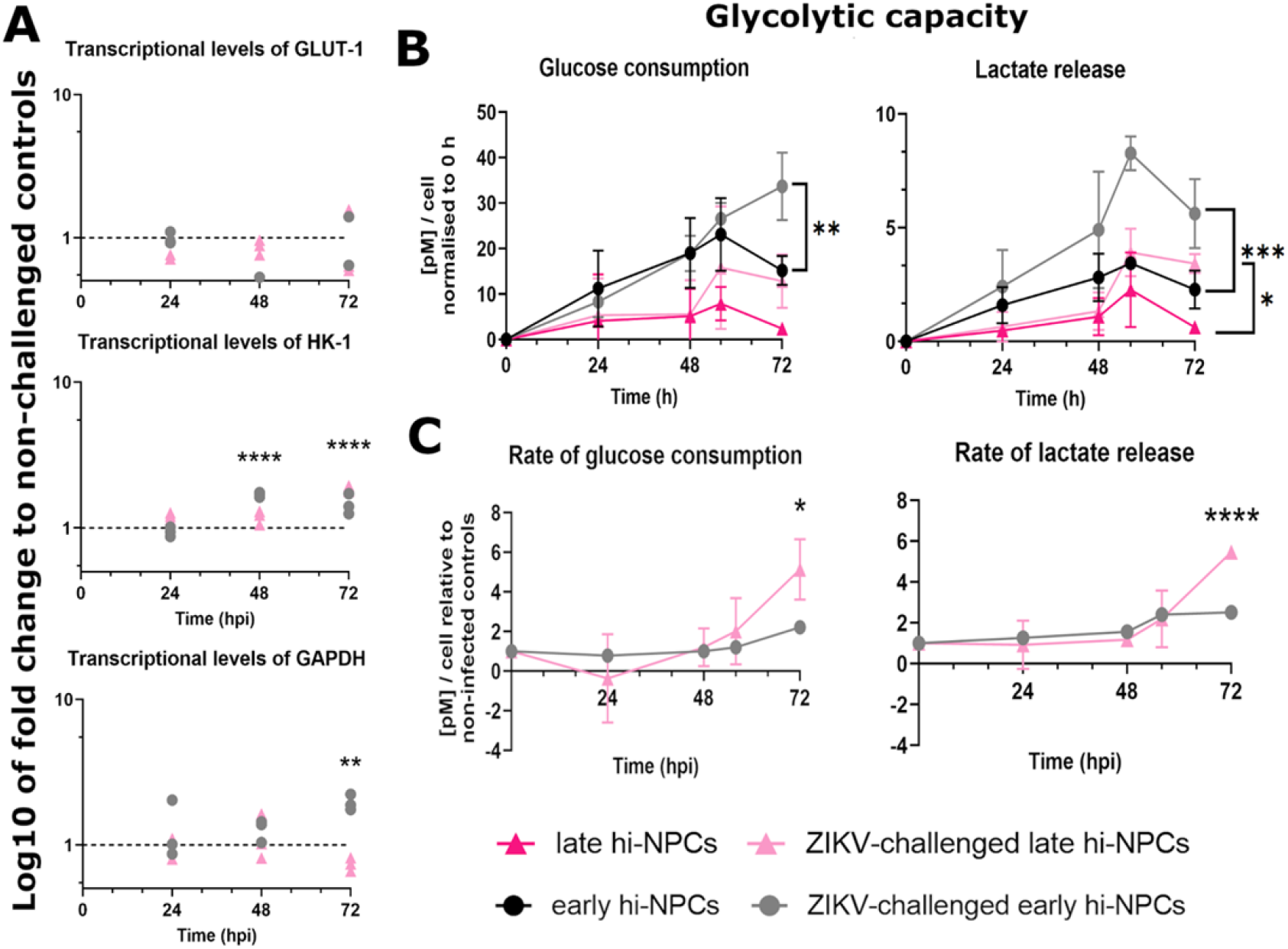
ZIKV induces a larger increment in the glycolytic capacity of late compared to early neuronal progenitors. The metabolism of glucose was assessed by (A) transcriptional levels of genes relevant for glycolysis in ZIKV-challenged hi-NPC subtypes relative to non-challenged controls. qPCR conducted in 4 replicates. Dots displaying the values of each donor cell line. Significance was calculated by Two-way ANOVA with Šidák’s multiple comparisons post-hoc test. Line graphs displaying (B) the calculated glucose consumption and lactate release from ZIKV-challenged and non-challenged hi-NPC subtypes and, (C) the ratio of change compared to their respective non-challenged controls. n = 2 viral challenges conducted in duplicates of three independent donors’ cell lines. Significance was calculated by mixed-effects model with Holm-Šidák correction. Error bars display mean ± SD. Significance is shown when *p < 0.05, ** p < 0.01, *** p < 0.001.

### ZIKV-mediated increase of cytosolic glucose metabolism is prominent in differentiating neuronal progenitors with lower glucose consumption

To assess whether ZIKV-induced differential alterations in hi-NPC subtypes were paralleled alterations in the metabolism of glucose, as a main substrate of neuronal progenitors, we quantified the transcripts for key glycolytic genes. Among these genes, HK-1 showed significant increases in ZIKV-challenged cells compared to non-challenged controls whilst no significant changes were calculated in the glucose entry receptor GLUT-1. Analysis of GAPDH showed a significant increased over time in early hi-NPCs whilst in late-hiNPCs significant increase was observed at early time points. Between neuronal progenitor subtypes, at 48 h.p.i., ZIKV-challenged early hi-NPCs showed a 1.42-fold increase in the gene expression of HK-1 compared to late hi-NPCs (Fig. 3A). However, at 72 h.p.i., a 1.26-fold increase was measured in the RNA levels of HK-1 in late hi-NPCs. At this time-point, early hi-NPCs showed a significant increase of 2.64-fold in the RNA levels of GAPDH compared to late hi-NPCs (Fig. 3A).

These effects were mirrored at the enzyme level with a significant increase in the overall glycolytic capacity in both progenitor subtypes. In ZIKV-challenged early hi-NPCs, there was a significant increase in the glucose consumption (p = 0.0053) at later stages of ZIKV replication whilst a greater lactate release (p ≤ 0.0101) was calculated in both hi-NPC subtypes when compared to non-challenged controls (Fig. 3B). As hi-NPC subtypes are metabolically distinct, we examined the possibility of ZIKV infection inducing a differential alteration in the metabolism of glucose by normalizing the glycolytic capacity in ZIKV-challenged hi-NPC subtypes to their respective controls. Data showed that, at 72 h.p.i., the increased glycolytic capacity induced during later stages of ZIKV replication was significantly different between progenitor subtypes. Late compared to early hi-NPCs showed a significantly increased the glucose consumption (p = 0.0122) and lactate release (p < 0.0001) with up to 2.31-fold and 2.17-fold changes, respectively (Fig. 3C).

### ZIKV-infected neuronal progenitor subtypes display specific patterns of mitochondrial alterations during viral replication

We examined the characteristics of mitochondrial homeostasis in ZIKV infected (Env +ve), ZIKV-challenged neighbouring cells (Env −ve), and non-challenged controls by immunostaining (Fig. 4A). Distinction between Env +ve and Env −ve was done by the immunodetection of viral proteins in challenged hi-NPCs. Quantification of different mitochondrial parameters by single-cell imaging using defined mitochondria areas of analysis (Fig. 4B) showed that cellular signalling from Env +ve cells, at 24 h.p.i., induced a significant increase in the mitochondrial membrane potential of Env −ve in early and late hi-NPCs subtypes (p = 0.045 and p = 0.0151, respectively) (Fig. 4C). We found that mitochondrial alterations in Env +ve hi-NPCs and non-challenged controls were distinct during stages of ZIKV replication and that these were specific to each neuronal progenitor subtype. Env +ve early hi-NPCs showed, at 24 h.p.i., a significant reduction in mitochondrial size (p = 0.0328) (Fig. 4D). From these cells, a significant reduction in mitochondrial abundance was also observed at 24 h.p.i. (p = 0.0091) and 48 h.p.i. (p = 0.0217) (Fig. 4E). Alterations in Env +ve late hi-NPCs showed that at 56 h.p.i., mitochondrial size (p = 0.0047) and abundance (p = 0.002) were significantly reduced (Fig. 4D and 4E). Lastly, mitochondrial distribution was significantly reduced at 24 h.p.i. in Env +ve early (p = 0.0251) and late hi-NPC subtypes (p = 0.0071) (Fig. 4F).

**Fig. 4.**
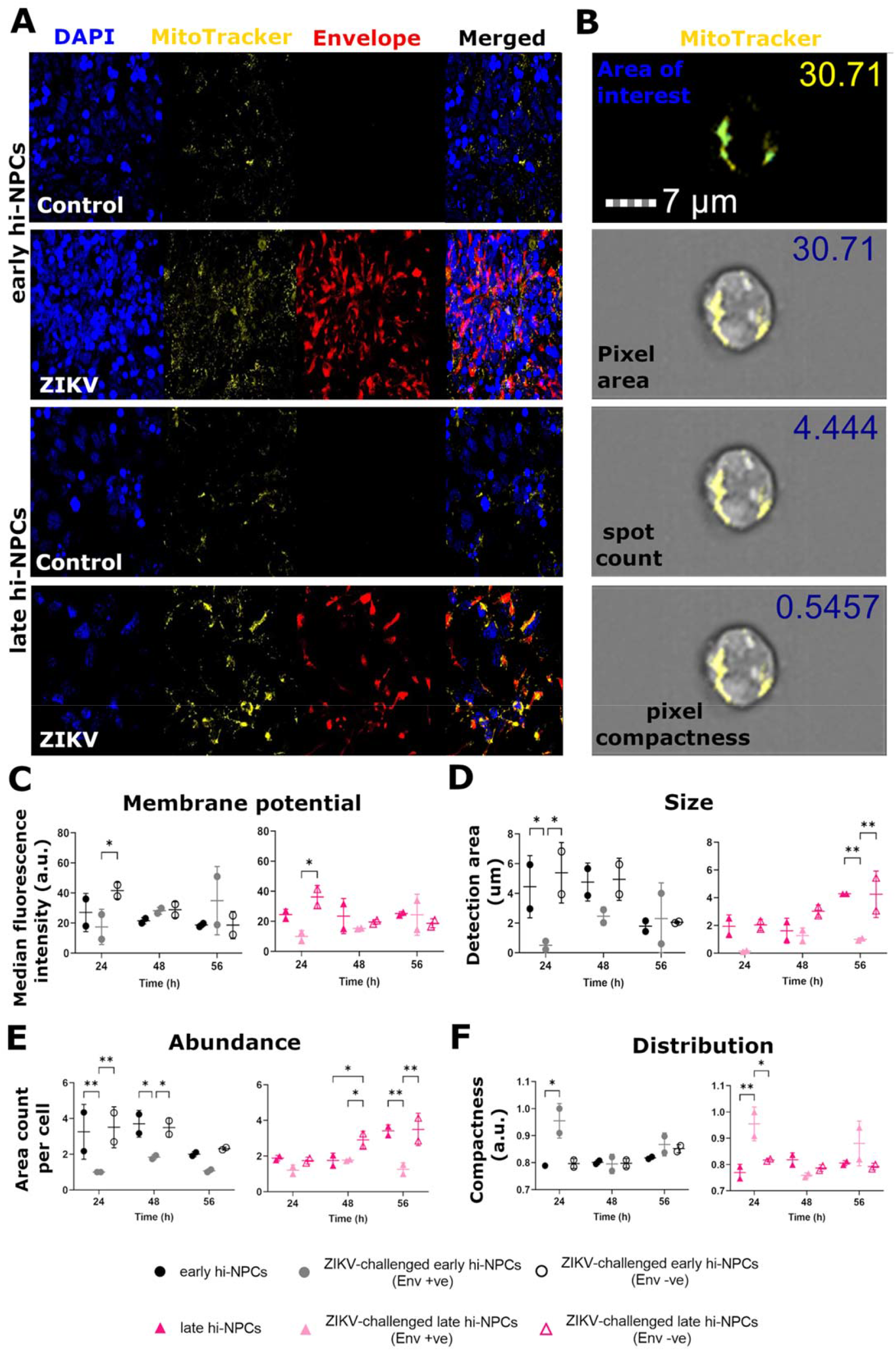
ZIKV induces neuronal progenitor subtype-specific mitochondrial stress. Analysis of several characteristics of mitochondria homeostasis in hi-NPC subtypes challenged to ZIKV compared to non-challenged controls. (A) Representative confocal images with high magnification (60x) of z-stack projections (15 μm) confirming positive staining of mitochondria (yellow) and ZIKV Envelope protein (red) in hi-NPC subtypes. DAPI staining (blue). (B) AMNIS Imaging flow cytometer digital images show representative mitochondria staining (yellow) with different quantified parameters, selected by defined mask (blue) in hi-NPC subtypes. Dot plots showing (C) mitochondrial membrane potential by measurements of the median fluorescence intensity of MitoTracker™ Red CMXRos, (D) mitochondria size estimated by the quantification of the total occupied area within the cells, (E) abundance of areas occupied by mitochondria and, (F) distribution of mitochondria determined by the proximity of positive areas within the cells. A minimum of 500 *in focus* cells were analyzed per donor cell line out of 10000 cells recorded. n = 1 viral challenge conducted in duplicates of two donors’ cell lines. Significance was calculated by mixed-effects model with Tukey’s correction. Error bars display mean ± SD. Significance is shown when *p < 0.05, ** p < 0.01, *** p < 0.001.

### ZIKV infection induces neuronal progenitor subtype-specific regulation of lipid metabolic genes

The presence of ZIKV likely alters the transcriptional profile of the host cells. We found that ZIKV simultaneously increased the expression of genes involved in of both beta-oxidation and lipid biosynthesis at different time-points during ZIKV replication (Fig. 5A). In addition, gene expression showed that infection, particularly at later stages (72 h.p.i.), increased the transcriptional profile but to different rates between hi-NPC subtypes (Fig. 5B and C). PDK2, essential in brain cells to promote fat oxidation(15), showed a significant increase (p = 0.0153) in early hi-NPCs compared to late hi-NPCs exclusively at 72 h.p.i. (Fig. 5B). The ACADM gene, required for the synthesis of enzymes for the oxidation of medium-chain fatty acids(16), showed at 24 h.p.i. a modest yet significant increase in late compared to early hi-NPCs (late:early RNA ratio of 1.27). In contrast, at 72 h.p.i., early hi-NPCs showed a significant 1.62-fold change in the levels of ACADM compared to late hi-NPCs (Fig. 5B). A significant increase in the HADHA gene (late:early RNA ratio of 1.26), essential for the synthesis of multi-enzymes within the mitochondrial trifunctional complex(17), was also observed exclusively at 24 h.p.i. (Fig. 5B). The analysis of genes involved in lipid biosynthesis(18) showed, only at 24 h.p.i., a significant differential expression of 1.64-fold in ACACA in late compared to early hi-NPCs. In contrast, the level of transcripts of ACACA at 48 h.p.i., showed a 1.33-fold change in early compared to late hi-NPCs (Fig. 5C). Transcriptional levels of FASN, the main biosynthetic enzyme involved in the synthesis of saturated long-chain fatty acids(18), showed similar results at 24 and 48 h.p.i. than those of ACACA. Late hi-NPCs showed a significant 1.43-fold increase compared to early hi-NPCs, exclusively at 24 h.p.i. whilst at 48 h.p.i., opposite results showed a modest 1.12-fold increase early compared to late hi-NPCs (Fig. 5C). Furthermore, single cell immunofluorescence analysis of lipid droplets in ZIKV-infected hi-NPCs (Fig. 5D) showed that genetic changes were not translated in alterations of the intracellular abundance of neutral lipids (Fig. 5E). In early hi-NPCs, exclusively at 56 h.p.i., Env +ve cells exhibited significantly reduced levels of lipid droplets compared to Env −ve cells (p = 0.0331) (Fig. 5E).

**Fig. 5.**
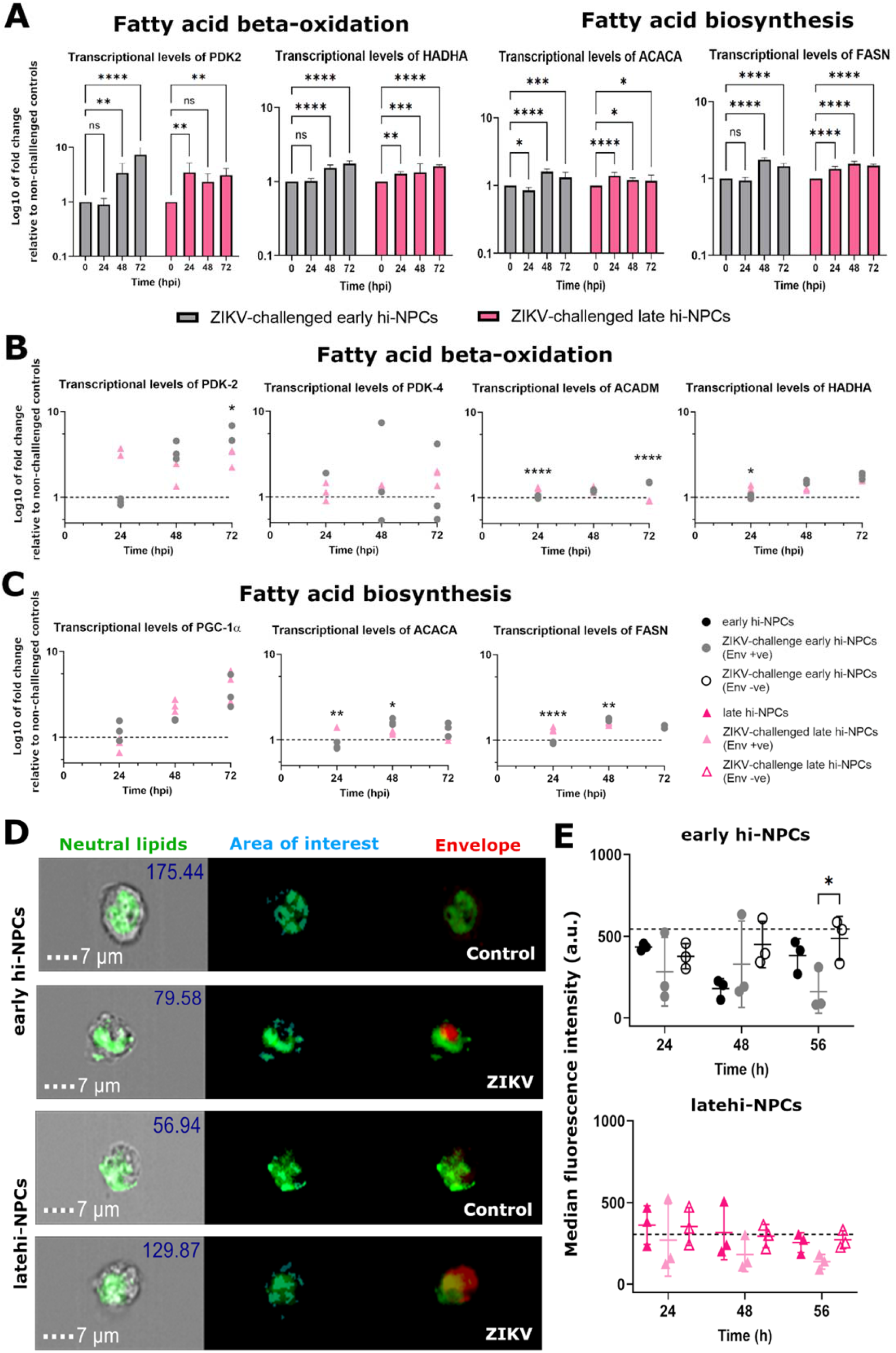
Course of ZIKV replication induces neuronal progenitor subtype-specific simultaneous differential gene expression of fatty acid beta-oxidation and biosynthesis. (A) Bar graphs showing the simultaneous upregulation of genes involved in fatty acid beta-oxidation and biosynthesis in neuronal progenitor subtypes during ZIKV replication. Bars displaying the 4 replicates. Dot plots showing the transcriptional levels of genes relevant in (B) fatty acid beta-oxidation and, (C) fatty acid biosynthesis in ZIKV-challenged hi-NPCs relative to non-challenged controls. qPCR conducted in 4 replicates. Dots displaying the values of each donor cell line. (D) AMNIS Imaging flow cytometer digital images showing representative lipid droplet staining (green), selected by defined mask (light blue), and ZIKV-Envelope staining (red). (E) Dot plots showing the abundance of lipid droplets in ZIKV Envelope positive, Envelope negative, and non-challenged hi-NPC controls. Values reflect the median fluorescence intensity (MFI) of CellTracker™ green BODIPY dye. A minimum of 500 *in focus* cells were analyzed per donor cell line out of 10000 cells recorded. n = 1 viral challenge conducted in three independent donors’ cell lines. Significance was calculated by Two-way ANOVA with Šidák’s multiple comparisons post-hoc test. Error bars display mean ± SD. Significance is shown when *p < 0.05, ** p < 0.01, *** p < 0.001.

### Initial stages of ZIKV replication dysregulate lipid droplet homeostasis exclusively in early neuronal progenitors

Lipid droplets, important during neurogenesis, are also crucial for the assembly of ZIKV particles(19). Thus, we examined whether there is a potential neuronal progenitor subtype specific alteration of lipid droplet homeostasis during ZIKV replication that may contribute to the disparities in brain damage at different trimesters. We analysed the intracellular lipid droplet content by immunostaining (Fig. 6A) and single-cell imaging (Fig. 5D). Early Env +ve hi-NPCs exhibited a significantly higher area occupied by lipid droplets (p ≤ 0.0242) compared to Env −ve and non-challenged control cells, followed by a time-dependent reduction (Fig. 6B). Data corresponding to the abundance of lipid droplets showed similar results with a significant increase (p ≤ 0.0189) in early Env +ve hi-NPCs compared to Env −ve and non-challenged controls (Fig. 6C). Late hi-NPCs showed no alterations during ZIKV replication (Fig. 6B and 6C). Distribution of the area occupied by lipid droplets showed a time-dependent increase in Env +ve hi-NPCs with significant differences at 56 h.p.i. observed exclusively between early Env +ve and Env −ve hi-NPCs (Fig. 6D). These data showed dysregulation of lipid droplets metabolism at early stages of ZIKV replication exclusively in early hi-NPCs yet, we did not find significant differences in the ratio of changes in lipid droplet homeostasis between ZIKV-challenged hi-NPC subtypes.

**Fig. 6.**
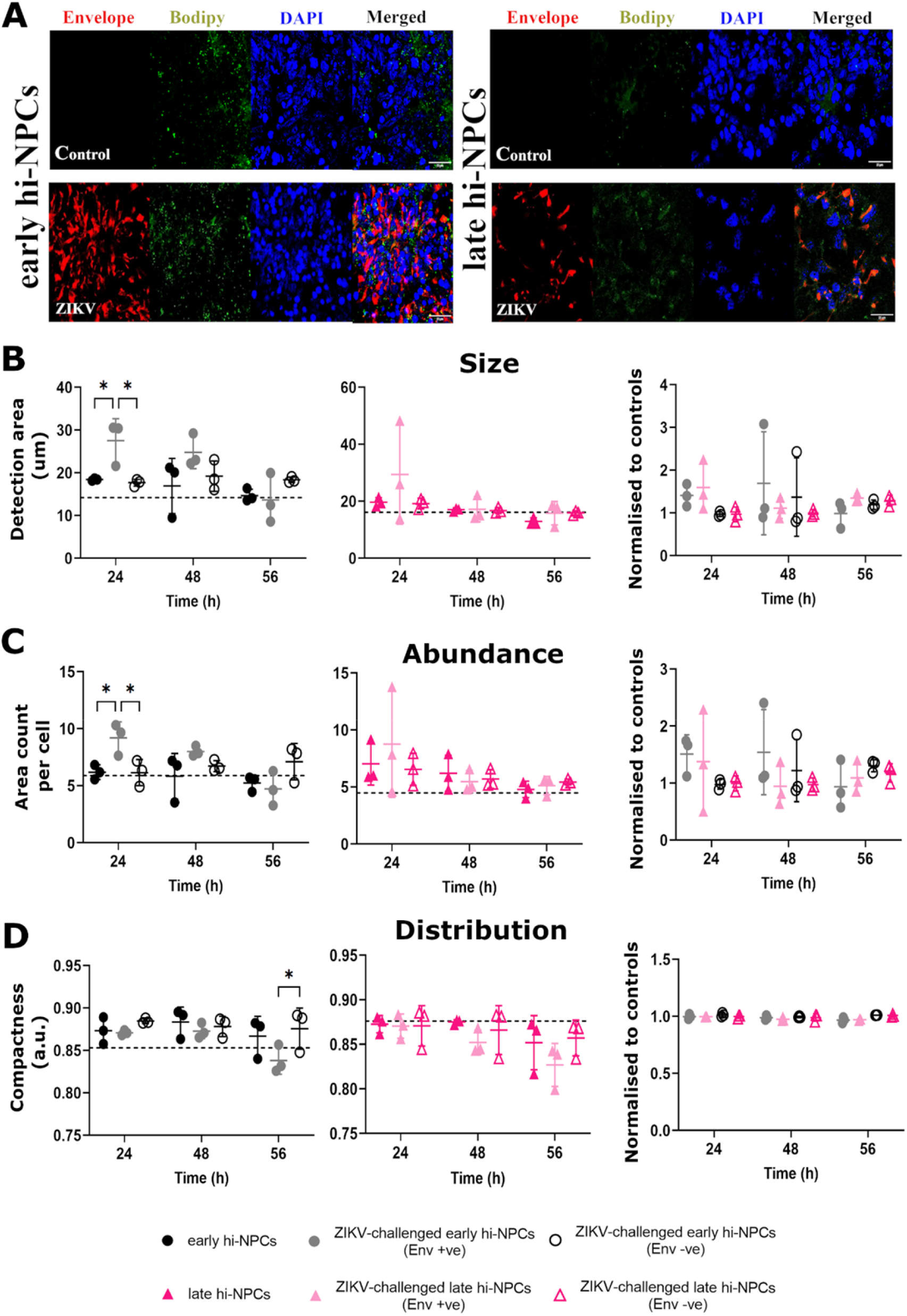
Early stages of ZIKV replication promotes accumulation of lipid droplets exclusively in early neuronal progenitors. Analysis of several characteristics of the lipid droplets in hi-NPC subtypes challenged to ZIKV compared to non-challenged controls. (A) Representative confocal images with high magnification (60x) of z-stack projections (15 μm) confirming positive staining of lipid droplets (green) and ZIKV Envelope protein (red) in hi-NPC subtypes. DAPI staining (blue). Dot plots showing (B) lipid droplet sizes estimated by the quantification of the total occupied area within the cells, (C) abundance of areas occupied by lipid droplets and, (D) distribution of lipid droplets determined by the proximity of positive areas within the cells. Graphs showing (left) the quantification by hi-NPC subtypes and (right) the ratio of change of each hi-NPC subtype relative to their respective non-challenge controls. A minimum of 500 *in focus* cells were analyzed per donor cell line out of 10000 cells recorded. n = 1 viral challenge conducted in three independent donors’ cell lines. Significance was calculated by Two-way ANOVA with Šidák’s multiple comparisons post-hoc test. Error bars display mean ± SD. Significance is shown when *p < 0.05, ** p < 0.01, *** p < 0.001.

## Discussion

3D neurospheres or organoids, differentiated from either fetal or iPSC-derived cells *in vitro*(20) are used to study fetal brain development. However, these systems show high variability, and this complicates the interpretation of differential responses to stimuli, including viral infection(21). Thus, to better understand the effects of ZIKV infection in different types of neuronal progenitors, we performed a detailed characterization of the populations of neuronal progenitors obtained from hiPSC (hi-NPC) using a modified existing 2D protocol(12). hi-NPCs showed two morphologically distinct cell populations at different time-points during differentiation. Both populations expressed generic cellular markers of neuronal progenitors and proliferation similar to that reported by others(12,13). However, significantly higher expression of Pax6 in late compared to early hi-NPCs may be correlated with Pax6 dynamic expression during neuronal proliferation and differentiation(22). This potential distinction of neuronal progenitor subtypes between cultures was supported by the significant differences in cellular metabolism.

The metabolic shift from the cytosolic metabolism of glucose to the mitochondrial oxidation of glucose-derived pyruvate and fatty acids is required to generate the pool of metabolites necessary to support neuronal differentiation(3–5). The metabolic profile of hi-NPC cultures showed contrasting results at each stage of differentiation. Early hi-NPC showed high glycolytic flux and low fatty acid oxidation, and late hi-NPCs showed a significantly decreased glycolytic flux and increased rate of fatty acid oxidation. These results, compared to human and mice neuronal progenitor subtypes(3–5,14), suggest that late hi-NPCs metabolically represent neuronal progenitors transitioning between quiescent radial-glia and proliferating precursors. In contrast, the metabolic profile of early hi-NPCs suggests these cells are likely to be quiescent radial-glia like cells. This was supported by the long-term estimation of the glycolytic capacity that, although it did not differ between hi-NPC subtypes, showed early hi-NPCs consumed higher levels of glucose, potentially due to differences in the length of their cell cycle. Thus, early and late hi-NPCs may recapitulate distinct populations of neuronal progenitors which their abundance changes over the course of forebrain development.

ZIKV successfully infects *in vitro* a pool of neuronal progenitors(8,9) yet, whether this infectivity is specific to a particular subtype remains unclear. We showed for the first time that ZIKV productively infects subtypes of neuronal progenitors yet to a different extent. Late hi-NPCs accumulate more ZIKV transcripts that result in a greater release of infectious ZIKV particles exclusively at 56 h.p.i. with the decay at later time-points potentially correlated with larger cellular death and thermal decay of previously released virions. Intracellular translation of viral RNA proteins causes several responses that may differentially contribute to the cell-type specific pathogenesis. Thus, no significant differences in the accumulation of intracellular envelope protein (E) between hi-NPCs may suggest that differential pathogenesis in these cells may be due cell-specific response viral hijacking. In accordance with different cellular models(23–25), both neuronal subtypes showed loss of cell viability after 48 h.p.i. Nuclear disruption in neuronal progenitors constitutes a feature of ZIKV replication, yet, this is the first report showing differential changes in nuclear morphology between neuronal progenitor subtypes. Our main finding was that viral perinuclear replication centres(26) (white arrows, Supplementary Figure 2) were only visible in late hi-NPCs and not in early hi-NPCs. Observed reduction in nuclear size and increased nuclear circularity in early hi-NPCs at specific time-points may be related to cellular shrinkage and death at the time of analysis(27).

*In vitro* passage of ZIKV may select for mutations that promote gain of infectivity, decreased cycle time and/or increased virus production(28), and these may in turn impact on cellular metabolism during replication. Thus, we used a low passage pathogenic African strain U-1962 (MP1751) to study the potential energetic requirements in neuronal progenitors. Hijacking of cellular metabolism by flaviviruses is a distinct process that varies between virus strains and host cells. For example, Dengue virus (DENV) infection directly increments and requires lactate glycolysis for its replication and, whilst this is not observed in West Nile virus (WNV), lipid biosynthesis is incremented during infection of both viruses; DENV significantly augments fatty acid biosynthesis and WNV cholesterol biosynthesis(8,9). Therefore, understanding whether ZIKV infection causes specific metabolic changes among different subtypes of neuronal progenitors or if it recapitulates metabolic features observed in other flaviviruses, potentially highlight mechanisms relevant in the pathogenesis of ZIKV in the developing fetal brain.

ZIKV infection resulted in transcriptional upregulation of genes involved in glycolysis in hi-NPC. Increased glucose consumption during ZIKV replication has previously been seen using a diversity of human and non-human cell lines(29–32). Increased glycose consumption was reflected by cytosolic processing of glucose and elevated lactate release during infection, as observed by others(21,30,31). Higher activation of glycolysis in late hi-NPCs, the subtype with lower baseline consumption rate of glucose, may suggest that ZIKV requires increased by-products of glycolysis to support additional processes such as redox clearance and nucleotide production(21), and/or to signal cell survival-related pathways necessary for efficient viral replication(33,34).

The role of mitochondria in the orchestration of metabolic pathways is likely to be manipulated by ZIKV(35). We showed that ZIKV-infected (Env +ve) hi-NPC have dysregulated mitochondrial homeostasis in the form of reduced size and number when compared to Env −ve and non-challenged controls. Interestingly, mitochondrial dysregulation showed to be time-dependent and differed between hi-NPC subtypes. This could be potentially explained by the specific metabolic characteristics of each neuronal progenitor subtype. Reduced distribution of mitochondria was found in both subtypes of hi-NPCs exclusively at the initial time-point, before quantifiable released of ZIKV particles had begun, potentially suggesting adaptative mechanisms to ZIKV replication. We also showed reduced mitochondrial membrane potential in Env +ve compared to Env −ve hi-NPCs but not to non-infected controls. These findings are different to those in ZIKV-infected human neuronal stem cells, human retinal and human hepatoma cells(36) in which ZIKV-infected cells had higher mitochondrial membrane potential.

Altogether, these results showed for the first time that ZIKV-induced dysregulation of mitochondrial homeostasis differs between neuronal progenitor subtypes and that mitochondrial changes, at specific time-points during viral replication, are exclusively observed in infected and not in neighbouring cells. Thus, highlighting the importance of time-course mitochondrial tracing to better understand the role of energy metabolism in the pathogenesis of ZIKV.

Results from human monocytes and drosophila suggest ZIKV infection increases beta-oxidation of fatty acids at early time-points(37,38). However, results obtained from mouse models of ZIKV pathology showing decreases in the TCA cycle, oxidative phosphorylation, and cytosolic levels of NAD^+^(30,33) may suggest a decreased beta-oxidation of fatty acids. We showed that ZIKV simultaneously increases the expression of genes involved in fatty acid beta-oxidation and biosynthesis yet to different rates between neuronal progenitor subtypes. Early hi-NPCs transitioned from increased beta-oxidation at early time-points to increased lipid biosynthesis whilst late hi-NPCs showed opposite patterns. The complex alterations of transcription of metabolic genes during ZIKV infection in hi-NPC subtypes may better reflect the situation in vivo than current cellular models and may enable one to resolve the current disparities in the literature(30,33,37,38).

Lipid droplets are multifunctional organelles that comprise aggregates of several types of lipids that can be exploited by ZIKV to support its replication(19) yet their entire role during ZIKV infection remains unclear. Changes in lipid droplet homeostasis during the course of infection have been showed in less relevant non-neuronal models(39). Thus, we examined the homeostasis of lipid droplets by their number and size during ZIKV infection. Our results showed that differential effects in the lipid droplet content between neuronal progenitor subtypes were restricted to ZIKV Env +ve early hi-NPCs. The significant increase in the lipid droplet abundance in Env +ve compared with Env −ve cells at 24 h.p.i. are in contradiction with the findings from a study of a similar kind in placental stromal cells that showed significant increases in lipid droplets in Env −ve cells(40). This potentially reflecting differences in the cellular response to ZIKV infection between cell types and/or the dynamics of lipid droplet utilization during sustained ZIKV infection.

To conclude, we showed that neuronal progenitor subtypes differentiated from hiPSC can be distinguished by their metabolic profile of glucose and fatty acids and that their metabolic differences are distinctively hijacked by ZIKV to support sustained replication. ZIKV replication induces an increase in the glycolytic capacity in all neuronal progenitor subtypes yet, mitochondrial and lipid droplet stress is specific to particular subtype (Fig. 7). This highlights a primary role of energy metabolism in neuronal progenitor subtypes of developing fetal brain in response to ZIKV infection that may potentially contribute to the distinct brain damage observed in new-borns when maternal infection occurs at different trimesters during pregnancy.

**Fig. 7.**
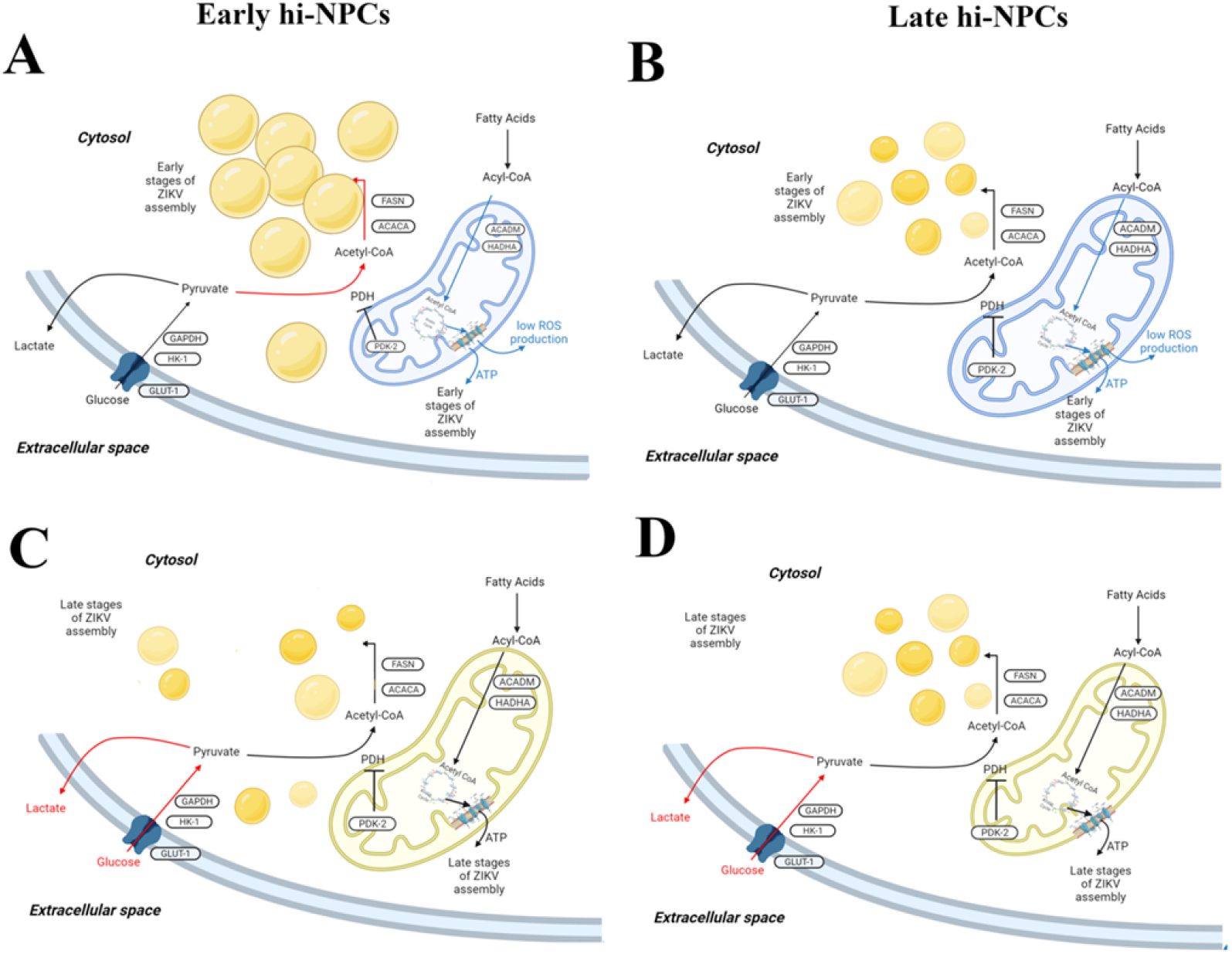
ZIKV-induced neuronal progenitor subtype-specific metabolic alterations. Schematic representation of the differential metabolic dysregulation between hi-NPC subtypes in response to ZIKV infection. (A) Increased number and area occupied by lipid droplets with reduced mitochondria membrane potential and size in early hi-NPCs during early stages of ZIKV replication. (B) Normal abundance of lipid droplets, normal mitochondria size and reduced membrane potential in late hi-NPCs during early stages of ZIKV replication. (C) Normal abundance of lipid droplets with larger distribution and increased glycolytic capacity in early hi-NPCs during late stages of ZIKV replication. (D) Normal abundance of lipid droplets, reduced mitochondria size and increased glycolytic capacity in late hi-NPCs during late stages of ZIKV-replication. Black arrows representing normal levels. Red arrows highlighting increased pathways. Light blue representing low mitochondrial activity with light yellow representing normal levels. Lipid droplet abundance and mitochondria size are represented within each figure. Pathways including some relevant metabolic genes.

## Methods

### Propagation and titration of ZIKV stocks

ZIKV strain (MP1751(41,42)) was propagated on Vero cells (ATCC CCL-81) passages 56 to 62. Cells were firstly incubated with the viral inoculum for 1.3 h at room temperature (RT) followed by further four days at 37 °C, 5% carbon dioxide until supernatant was collected upon evident cytopathic effect (CPE). Virus titration conducted by plaque assay was done by serial 10^−1^ viral dilutions in Vero cells at a density of 2.5 × 10^5^ cells. 1.5% carboxymethyl cellulose overlay was added after 2 h incubation to prevent viral spread. Plates were further incubated for 80 h under these conditions before plaques were reveal by the staining using Amido black for 30 minutes at RT. Plates were imaged using a Molecular Imaging ChemiDoc™ XRS+. Plaque forming units (p.f.u.) were calculated as follows: *p*. *f*. *u*. = (*n*/0.1)/*D* where *n* = average number of plaques, 0.1 is the volume of virus added in mL, and *D* = dilution factor.

### Viral thermostability

Thermostability of ZIKV at 37 °C was assessed by plaque assay. 500 μL of media-containing virus were added to wells of a 24-well plate, each corresponding to a specific time point. 350 μL of media were collected and frozen down at −80 °C before titration.

### Culture of Vero CCL-81 cells

Vero CCL-81 cells passaged every 2-3 days were cultured at 37 °C, 5% carbon dioxide in DMEM high glucose supplemented with 10% FBS. Cell passage was done at ~90% confluency. All cells were pelleted by centrifugation at 400g for 5 minutes at 4°C prior to resuspension. For the preparation for plaque assays, media from Vero cells was replaced for DMEM high glucose supplemented with 1% FBS and 1% Penicillin-Streptomycin (P/S).

### Human induced pluripotent stem cells (hiPSCs)

The hiPSC lines used in this study have been reported elsewhere: SFC840-03-03(43), SFC841-03-01(44) and, SFC856-03-04(45). Cell lines were derived from dermal fibroblasts from disease-free donors recruited through StemBANCC(46) and the Oxford Parkinson’s Disease Centre: participants were recruited to this study having given signed informed consent, which included derivation of hiPSC lines from skin biopsies (Ethics Committee: National Health Service, Health Research Authority, NRES Committee South Central, Berkshire, UK, who specifically approved this part of the study (REC 10/H0505/71)). Non-sendai reprogramming (Cytotune, Life Technologies) was used to reprogram fibroblast cells into hiPSCs. hiPSCs were cultured in defined, open-source medium termed OXE8(47). Cells, resuspended as clumps by using 0.5 mM EDTA, were plated onto Geltrex™ precoated plates and cultured at 37 °C, 5% carbon dioxide. Media changes were done every 24 h using not supplemented OXE8(47). At ~90% confluency, cells were passaged. Cells were passaged every 2-3 days and, after passage four, cells were either differentiated or stored in liquid nitrogen (LN_2_).

### Generation and culture of cortical progenitor cells (hi-NPCs)

hiPSC passaged 2-4 times post-thawing at a confluency of ~95% were induced to neuronal lineages using a modification of a protocol reported elsewhere(13). For neuronal differentiation, 6-7 days incubation in neuronal induction media (NIM) were required. Media was replaced daily and cells were observed under the microscope to assess the formation of a neuroepithelium-like sheet. If detachment (normally occurring after day 6), cells were washed once with PBS and incubated with 0.5mM EDTA in PBS for 5 minutes at 37 °C, 5% carbon dioxide. Cells were pelleted at 300 rcf for 3 min and resuspended, as clumps, in neuronal maintenance media (NMM) supplemented with 10μM ROCKi. Media was replaced daily with NMM. After 5 days, cells were replated (split 1:2) as detailed before. Following 3 extra days in culture, cells were either replated (split 1:2) or stored in LN_2_ (early hi-NPCs). Two additional passages, each of 3 days, were further conducted. Cells from the final passage were replated as single cells suspension (late hi-NPCs). For long term storage in LN_2_, cells were resuspended in freezing media composed of NMM supplemented with 10% DMSO and 5 μM ROCKi. See Extended Figure 4 for details on media composition and volumes of preparation.

### ZIKV challenge in monocultures

For viral challenge in hi-NPCs, cells were incubated for 2 h at 37 °C, 5% carbon dioxide with media-containing ZIKV at an M.O.I. of 1. Media-containing virus was then removed, and cells were washed once with PBS before adding fresh NMM and incubated under the same conditions. Mock challenges were done by exposing cells to supernatants of Vero cells cultured for 72 h.

### Confocal microscopy

For detection of neuronal markers, permeabilised fixed cells were incubated overnight with primary antibodies in PBS with 0.3% triton X-100 and 5% DS at 4 °C. Antibodies dilutions were as follow: anti-Pax6 (1:200, Biolegend; Poly19013), anti-Pax6 (1:200, GeneTex; GTX113241), anti-Tbr2 (1:50, AbCam; Ab23345), anti-Tuj1 (1:1000, Biolegend; 801202), anti-Sox2 (1:150, Merck; Ab5603), anti-Iba1 (1:500, AbCam; Ab5076), anti-MAP2 (1:100, Sigma; M4403), anti-Nestin (1:100, AbCam; Ab93666), anti-NeuN (1:150, Merck; MAB377) and, anti-pH-3 (1:50, Abcam; ab14955). Incubation with secondary antibodies (1:500; goat anti-mouse and goat anti-rabbit Alexa Fluor 488, Alexa Fluor 546, Alexa Fluor 555, Alexa Fluor 647 and, Alexa Fluor 750, or donkey anti-goat Alexa Fluor 488 and Alexa Fluor 647 (all from Invitrogen)) was conducted for 2 h at RT. Cells were incubated for 5 minutes with 1:10.000 dilution of DAPI and mounted on a glass slide. For staining of mitochondrial membrane potential, live hi-NPCs were incubated with 130 nM MitoTracker™ Red CMXRos for 30 minutes in the dark at 37 °C, 5% carbon dioxide before fixing with 2% PFA for 30 minutes. Cells were permeabilised and blocked in FACs buffer (PBS supplemented with 1% FBS, 10 μg/mL human-IgG, and 0.01% Sodium azide) 0.01% saponin for 30 minutes at RT. Cells were incubated with anti-ZIKV Envelope (1:50, GeneTex; GTX133314) overnight at 4 °C. Lipid droplet staining (1:5000 BODIPY™ 493/503) was then conducted for 1 h at RT together with 1:2000 DAPI and goat anti-rabbit Alexa Fluor 647. Cells were then mounted, and images were acquired with a Zeiss LSM710 confocal microscope. Acquisitions were performed in five random fields captured under 63x magnification, 1.2 zoom of six sections per image, each section of 5 μm. Images were processed and/or quantified using Fiji-ImageJ2 version 2.3.0/1.53f.

### Flow cytometry

Resuspended hi-NPCs were live stained (LIVE/DEAD™ Fixable Violet Dead Cell Stain Kit) following the manufacturer’s procedure. Cells were then fixed with 2% PFA in PBS for 15 minutes or live stained with markers of brain cell types prior to fixation. Staining of fixed cells was conducted by an initial permeabilization (0.3% saponin FACs) for 1 h at RT followed by an overnight incubation at 4 °C incubation in primary antibody solutions Primary antibodies: anti-Pax6 (1:150, GeneTex; GTX113241), anti-Tbr2 (1:100, AbCam; Ab23345), anti-Sox2 (1:70, Merck; Ab5603), anti-MAP2 (1:100, Sigma; M4403), anti-Nestin (1:70, AbCam; Ab93666), anti-NeuN (1:200, Merck; MAB377), anti-Ki67 (1:500, Abcam; ab16667) and anti-S100B (1:30, Sigma; S2532). Secondary staining consisted of 1:2000 goat anti-rabbit Alexa Fluor 555 and goat anti-mouse Alexa Fluor 647, both from Invitrogen. Live staining of markers of brain cell types was conducted following the manufacturer’s procedure (BD Biosciences Human Neural Cell Sorting Kit). Shortly, after live/dead staining, cells were filtered through a 70 μm cell strainer and resuspended in 5 mM EDTA. Cells were then stained with either primary conjugated antibodies or isotype controls for 30 minutes at 4 °C. Finally, cells were washed once and resuspended in 1% PFA in FACs buffer for 30 minutes. Data acquisition was done with Cytoflex LX (Beckman Coulter) CytExpert software. Data analysis was performed using FlowJo V10.8.1.

### Fluorescence imaging

For nuclear imaging, hi-NPCs plated on 96-well plates were fixed with 2% PFA for 15 minutes at RT and stained with 1:10.000 DAPI dilution in PBS for 10 minutes. Cells were imaged with an EVOSFL Auto. Nuclear size and shape, of at least 300 nuclei per condition, were calculated using Fiji-ImageJ2 version 2.3.0/1.53f.

### Imaging flow cytometry (Imagestream^®^)

Resuspended hi-NPCs were live stained for mitochondrial membrane potential, fixed and permeabilised as previously detailed. Overnight incubation at 4 °C with primary antibody (FACs buffer 0.1% saponin, 1:100 anti-NS4A (GeneTex; GTX133704) and 1:100 anti-NS1 (Abcam; ab214337) was conducted prior to secondary staining with 1:1000 goat anti-mouse Alexa Fluor 546 and goat anti-rabbit Alexa Fluor 750. Cells were then re-stained over 3 h at RT with FACs buffer 0.1% saponin, 1:100 anti-Envelope. Secondary goat anti-rabbit Alexa Fluor 647 staining was conducted in parallel with 1:10.000 Bodipy™ staining for 1 h at RT. 1:10.000 DAPI staining was then carried out for 5 minutes. Cells were kept in PBS and data acquisition was done with an Amnis^®^ ImageStream^®X^ MkII (Luminex) Inspire 10 software. Data analysis was performed using IDEAS software V6.2.

### Glycolytic flux

hi-NPCs pre-plated for 10 hours (5 h post-attachment) were fed with culture medium containing 5-^3^H-glucose and incubated at 37 °C, 5% carbon dioxide. 5-^3^H-glucose is converted via glycolysis into fructose-6-phosphate releasing ^3^H_2_O into the culture medium. Culture media was composed of no glucose DMEM supplemented with 5 mM glucose and 0.2 μCi/mL or 0.00074 MBq/mL of glucose, D-[5-^3^H(N)] (PerkinElmer). Cells were incubated for 4 and 6 h after which cell supernatants from duplicate plates were collected at each respective time-point and stored at −20 °C. Released ^3^H_2_O required to be separated from unconverted 5-^3^H-glucose. The separation process was done using the ion-exchange chromatography separation (Dowex) method. Dowex solution was prepared by mixing 250 g AmberChrom^®^ 1X4 chloride form, 100-200 mesh with 1.25 M NaOH and 1.61 M boric acid. The mixture was mixed gently and repeatedly washed with dH_2_O pH 7.5 was reached. Dowex soltion was added to glass Pasteur pipettes (VWR 612-1701, Avantor; 612-1701) containing glass wool. 200 μl of the media were added to the Dowex column and incubated for 15 mins allowing ^3^H glucose to bind to the column. ^3^H_2_O was eluted to the vials by two rinsing them twice with dH_2_O. Radioactivity was measured using Tri-Carb 2800TR Liquid Scintillation Analyzer (Perkin Elmer). 0.2 mL medium at time point 0 was used as a control to determine the specific activity of the buffer.

### Glucose oxidation

Glucose oxidation was measured using a protocol modification(48) of the original CO_2_ capture method by Collins et al., 1998(49). hi-NPCs pre-plated for 10 h in a 24-well plate were washed once with PBS and fed with no glucose DMEM supplemented with 12 mM glucose containing 0.2μCi/mL [D-[14C(U) glucose: 1mCi-37MBq]. The released ^14^CO_2_ was trapped in KOH-soaked filter papers. To kill the cells and allow the released of ^14^CO_2_, cells were incubated with perchloric acid for 1 h after 2 and 4 h post-feeding. Filter paper containing trapped ^14^CO_2_ were analyzed using a scintillation counter and the radiation counts per minute (CPM) were measured. 0.01 mL medium at time point 0 was used as a control to determine the specific activity of the buffer.

### Oleic acid oxidation

Radioactive tracing of oleic acid oxidation was based on the detection of ^3^H_2_O produced from ^3^H oleate in the electron transport chain. hi-NPCs pre-plated for 10 h were fed for 4 and 6 h at 37 °C, 5% carbon dioxide with no glucose DMEM, 2% BSA, 0.3 mM oleate and 0.2μCi/mL of oleate, [9,10-3H(N): 1mCi-37MBq] (PerkinElmer). For media preparations, oleate was heated to melt them down and, was simultaneously added with ^3^H oleate to the BSA to ensure the same binding. Folch extraction method, modified and reported by Malandraki-Miller, S. et al., 2019(50) was used to separate the ^3^H_2_O from the ^3^H oleate. 0.5 mL of perfusate was pipetted in 15 mL falcon tube containing 1.88 mL of chloroform:methanol (1:2 v/v) solution, 625 μl chloroform and 625 μl KCL-HCl solution (2 M KCl, 0.4 M HCl). The solution was then rotated on a laboratory stuart rotator SB3 at 40 rpm for 1 h. After rotation, top aqueous layer was collected, and bottom organic layer was discarded. The aqueous layer was then exposed to 1 mL chloroform, 1 mL methanol and 0.9 mL KCL-HCL and rotated for 1 h at 40 rpm. 0.5 mL the top aqueous layer was used to count the radioactivity using a Tri-Carb 2800TR Liquid Scintillation Analyzer (Perkin Elmer). 0.5 mL medium at time point 0 was used as a control to determine the specific activity of the buffer.

### Extracellular glucose measurements

The concentration of glucose in the culture media was detected using a miniaturization of a commercial protocol (Merck; GAHK20). 100 μL of culture media were diluted 1:10 and 1:20 in dH_2_O. 20 μL of each dilution were added per well on a F-bottom 96-well plate. Glucose solution was freshly made up and 100ul were added onto each sample. Plates were incubated at 35 °C for 15 min inside a SpectraMax M5 plate reader. NADH absorbance was read at 340 nm wavelength. Glucose values from the samples were calculated by curve fitting to a known standard curve.

### Extracellular lactate measurements

Lactate from the culture media was based on the NADH absorbance at 340 nm wavelength. 5μL of sample were added per well and incubated with 100 μL of freshly made running buffer solution for 20 minutes at 37 °C inside a SpectraMax M5 plate reader. NADH absorbance was read at 340 nm wavelength. Lactate values were calculated by curve fitting to a known standard curve using sodium L-lactate.

### Cell survival measurements

CCK-8 assay was used to determine cell survival following manufacturers’ procedure. Cells were washed once with PBS and incubated for 2 h at 37 °C, 5% carbon dioxide with 100 μL fresh medium containing 10 μL of CCK-8. After incubation, 100 μL were collected and absorbance was read at 460 nm wavelength using a SpectraMax M5 plate reader.

### RNA extraction and cDNA conversion

Cells were detached with accutase and pellets were frozen dry at −80 °C. Cell pellets were thawed on ice and RNA extraction was done using the RNeasy Mini Kit. Samples were DNAse treated RNAse-Free DNase Set. RNA concentration was quantified using a Nanodrop 2000c. RNA was reversed transcribed into complementary DNA (cDNA) using the High-Capacity RNA-to-cDNA™ Kit. For cDNA conversion, samples were diluted, and equal concentration of RNA was added for the reaction.

### Quantitative polymerase chain reaction

Gene expression was determined using the double stranded DNA binding dye SYBR^™^ Green. Primers efficiency of designed non-published primers was determined by SYBR green qPCR across 5×3-fold dilutions of cDNA and calculated in Microsoft Excel as *Efficiency* = (10^(−1/*Slope*) − 1) * 100 where slope is calculated for the plot of average Ct against Log(sample quantity) – see Extended Data table 2. qPCR reactions comprised of 1 volume of sample cDNA to 3 volumes mastermix (Power SYBR^™^Green PCR Master Mix, forward and reverse primer mix, and nuclease free water at a ratio of 5μL : 1μL : 1.5μL respectively). Reactions were run in five replicates in either 96-well format, 20 μL/sample, or 384-well format, 10 μL/sample. The threshold cycle (2^−ΔΔCt^) method of comparative PCR was used to analyze the results. 2^−ΔCt^ was calculated by the normalization of the sample Ct value to the average Ct value of two housekeeper genes, UBC and TBP. 2^−ΔΔCt^ analysis was done by the further normalisation of the samples against a 0 h control. For the quantification of ZIKV transcripts, Ct values were also normalized against UBC and TBP and further normalised against unchallenged controls, displaying results as 2^−ΔCt^ and 2^−ΔΔCt^.

### Statistical analysis and software

Datasets from single experiments with large sample size, to compute for the time-course comparisons, were analysed by non-parametric paired Wilcoxon tests. Datasets containing missing values and/or values of zero after normalization from independent experiments of three donors’ cell lines were analyzed using mixed-effects model with Holm-Šidák correction. Datasets comprising two independent repeats of a donor cell line were analyzed by two-tailed Mann-Whitney U test. Datasets containing analysis of two independent donors or independent triplicate experiments of one donor cell line were analyzed using mixed-effects model with Šidák correction. Datasets comprising one repeat of two donors’ cell lines were analyzed by mixed-effects model with Tukey’s multiple comparisons test. Datasets containing analysis of independent experiments of three donors’ cell lines comprising two groups were processed using Two-way ANOVA with Šidák’s multiple comparisons post-hoc test whilst when comprising three or more groups post-hoc were computed using Tukey’s multiple comparisons test. All the statistical tests were performed using GraphPad Prism 9.3.1. A calculated P value less than 0.05 was reported as significantly different. Representation schemes were created with BioRender.com. Figures were done using GNU Image Manipulation Program (GIMP 2.8.22).

## Acknowledgments

Strain MP1551 was shared by the National Infection Service at Public Health England. The authors thank Dr. Sally Cowley, Dr. Sharat Warrier, Dr. Matthew Kerr, Dr. Robert Hedley, Dr. Maeva Dupont, Ms. Catriona Rooney and Ms. Vicky Ball (University of Oxford) for technical assistance. Also, the authors would like to thank the Flow cytometry facility at the Sir Dunn School of Pathology, the Cardiac Metabolic Research Centre – CMRG, the Molnár lab and the James and Lillian Martin Centre (University of Oxford). This work was supported by the James and Lillian Martin Centre. J.G.-J. is supported by Ecuadorian National Government Scholarship (Senescyt – Secretaria Nacional de Educacion Superior, Ciencia y Tecnologia). U.P. is supported by Indonesia Endowment Fund for Education (LPDP).

## Author contributions

J.G.-J. conceptualised the project. W.S.J. supervised the work. J.G.-J. performed the work and analysed the data. J.G.-J. designed the radioactive work which was conducted by U.P.. W.S.J. and Z.M. provided intellectual expertise. J.G.-J. wrote the original draft. W.S.J. and Z.M. reviewed the manuscript.

## Extended Figures

**Extended Fig. 1.**
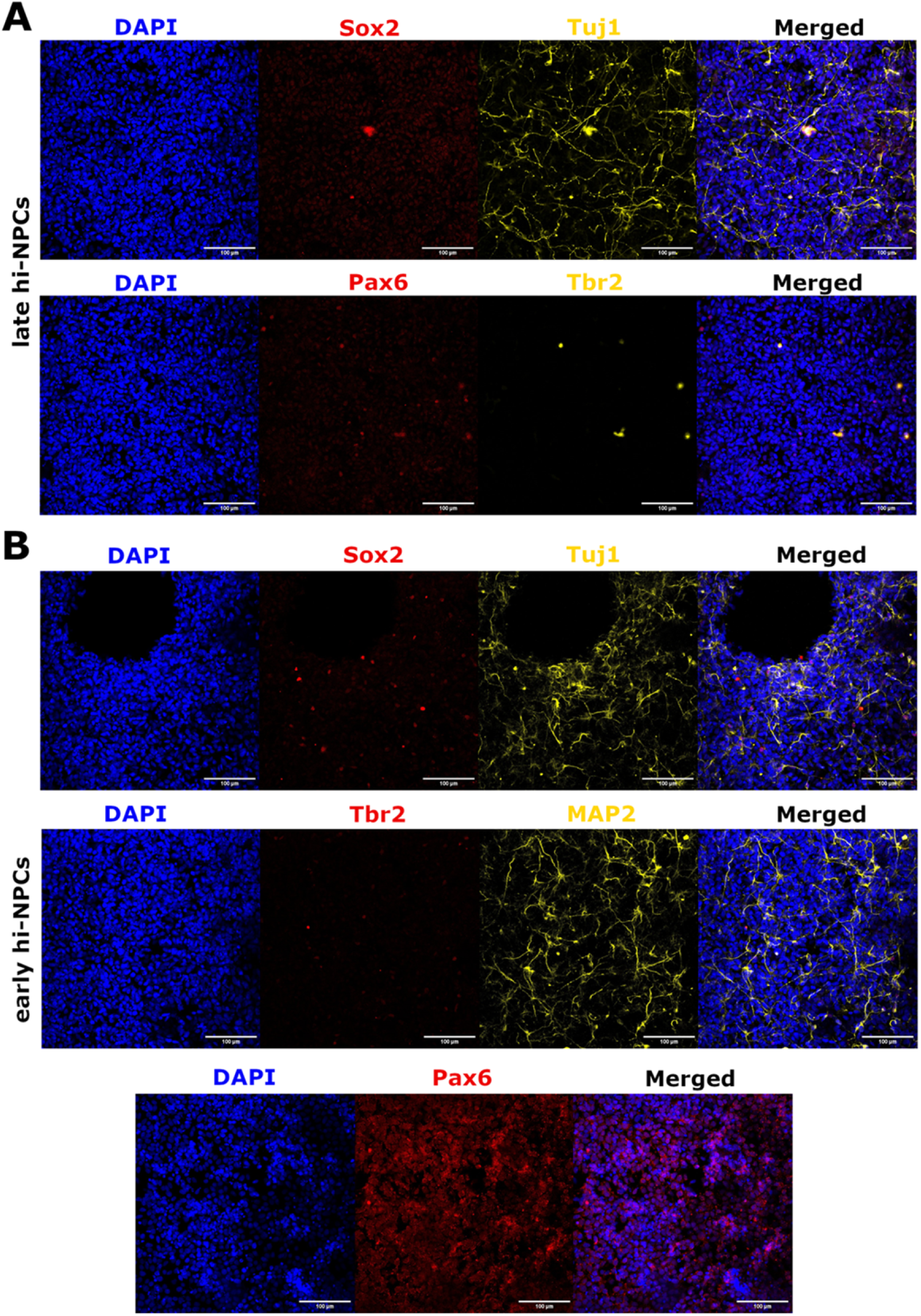
Phenotypically distinct cortical neuronal progenitors differentiated from hiPSCs express markers of progenitors of the forebrain. Representative confocal images (10x) of the detection of markers of *in vitro* neuronal progenitors of the forebrain (Sox2, Tuj1, Tbr2, Pax6 and MAP2) in early and late hi-NPCs. Scale bar: 100 μm.

**Extended Fig. 2.**
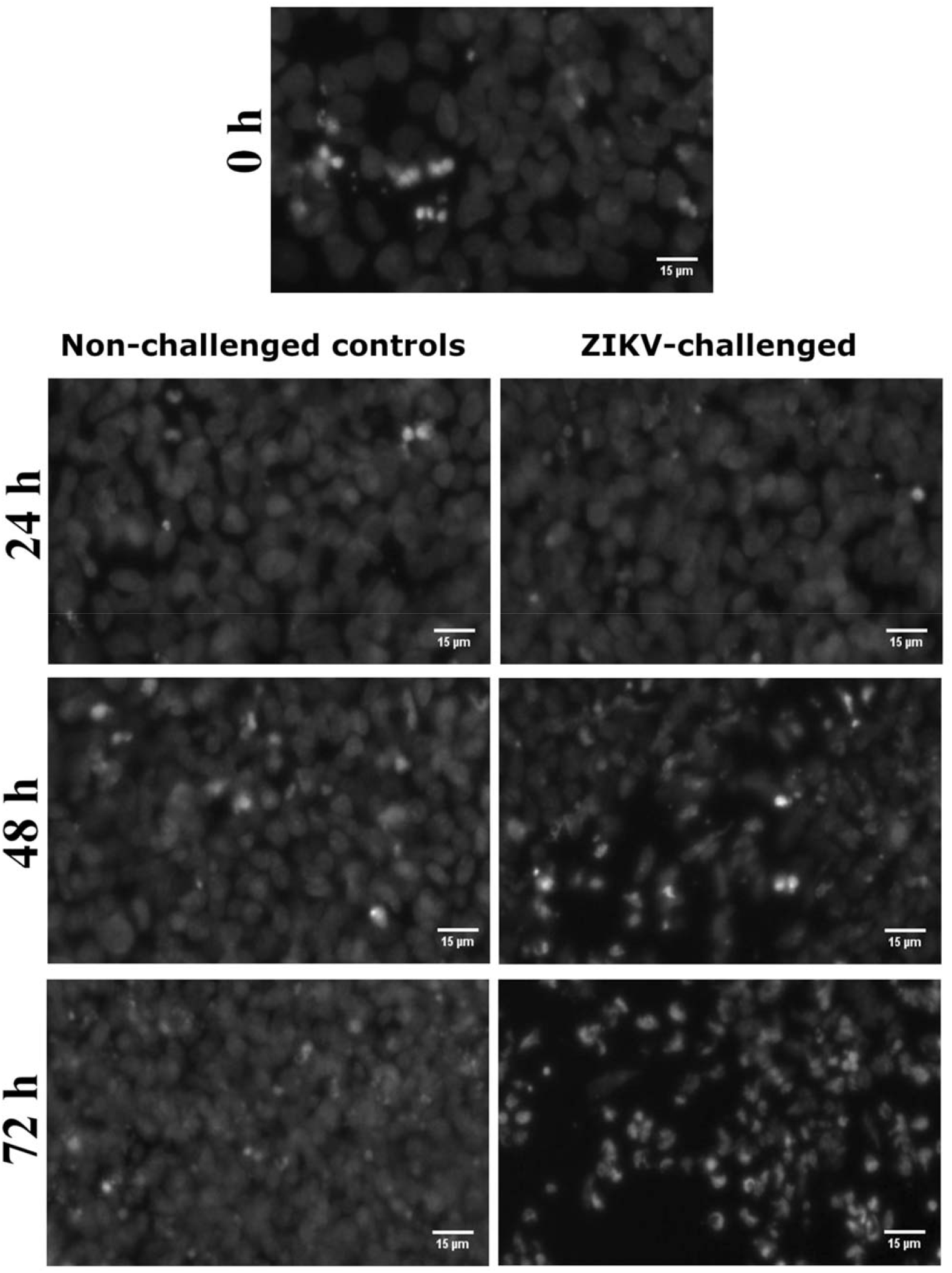
ZIKV infection causes nuclear damage in early hi-NPCs. Imaging (40x) of the nuclei of early hi-NPCs challenged to ZIKV over the course of 72 hours and the respective non-challenged controls. Nuclei were stained with DAPI.

**Extended Fig. 3.**
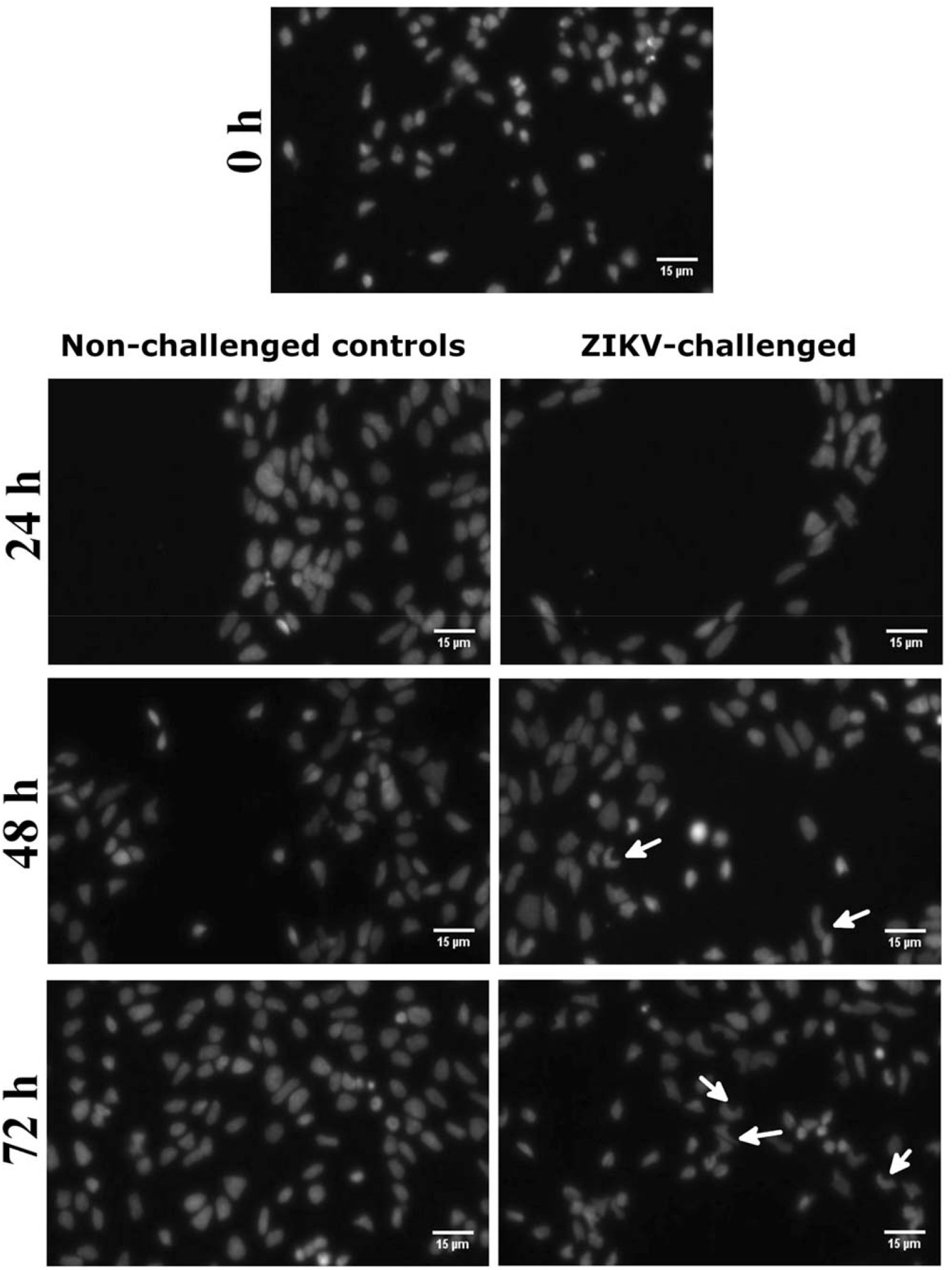
ZIKV show perinuclear replication sites in infected late hi-NPCs. Imaging (40x) of the nuclei of late hi-NPCs challenged to ZIKV over the course of 72 hours and the respective non-challenged controls. Nuclei were stained with DAPI. White arrows highlighting potential sites of viral factories.

**Extended Fig. 4.**
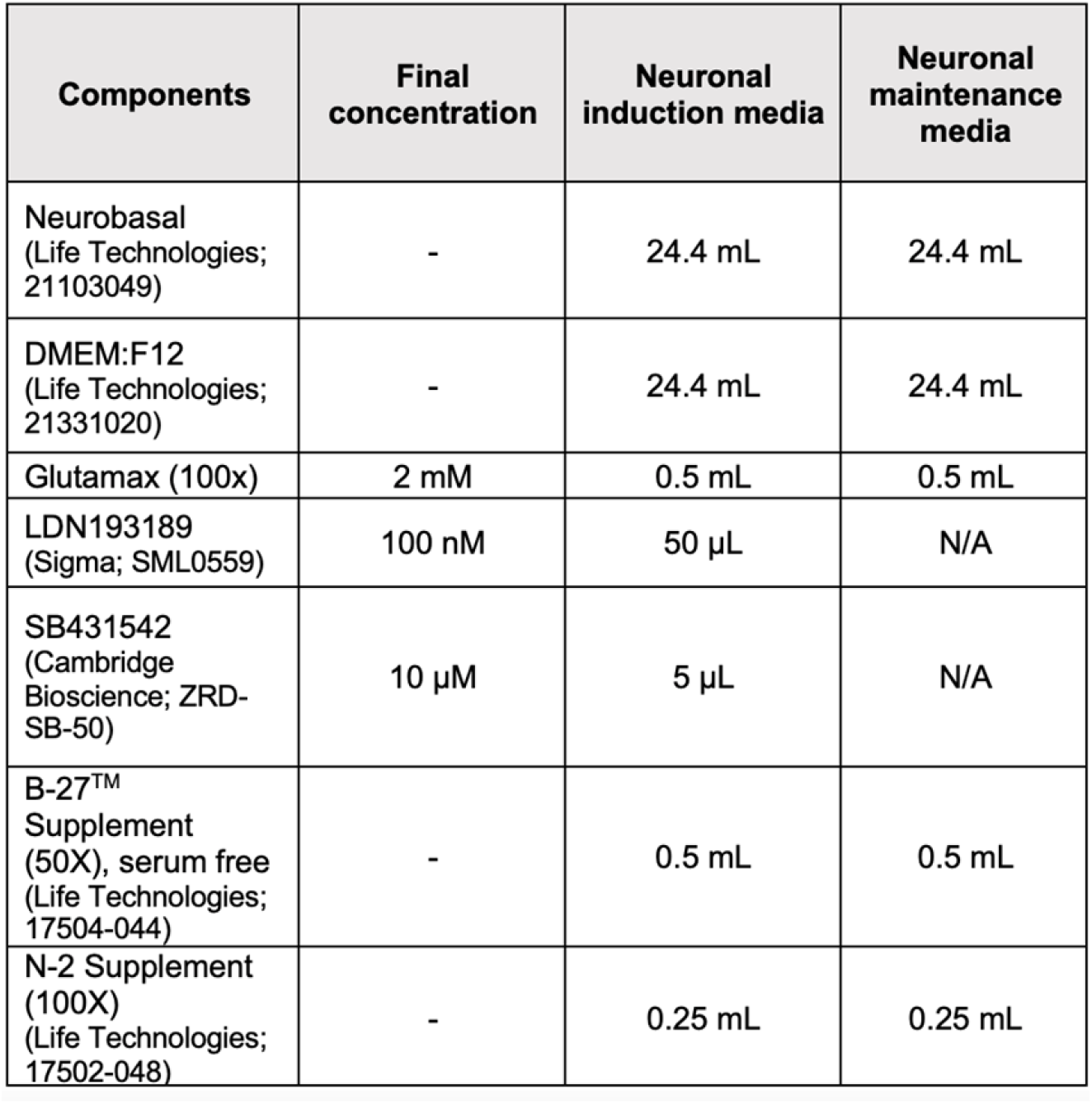
Media culture recipe for neuronal differentiation from hiPSC. Figure showing the volumes and concentration of the reagents used to differentiate cortical neuronal progenitors from hiPSC. The volumes represented on the table are to prepare 50 mL.

**Extended Fig. 5.**
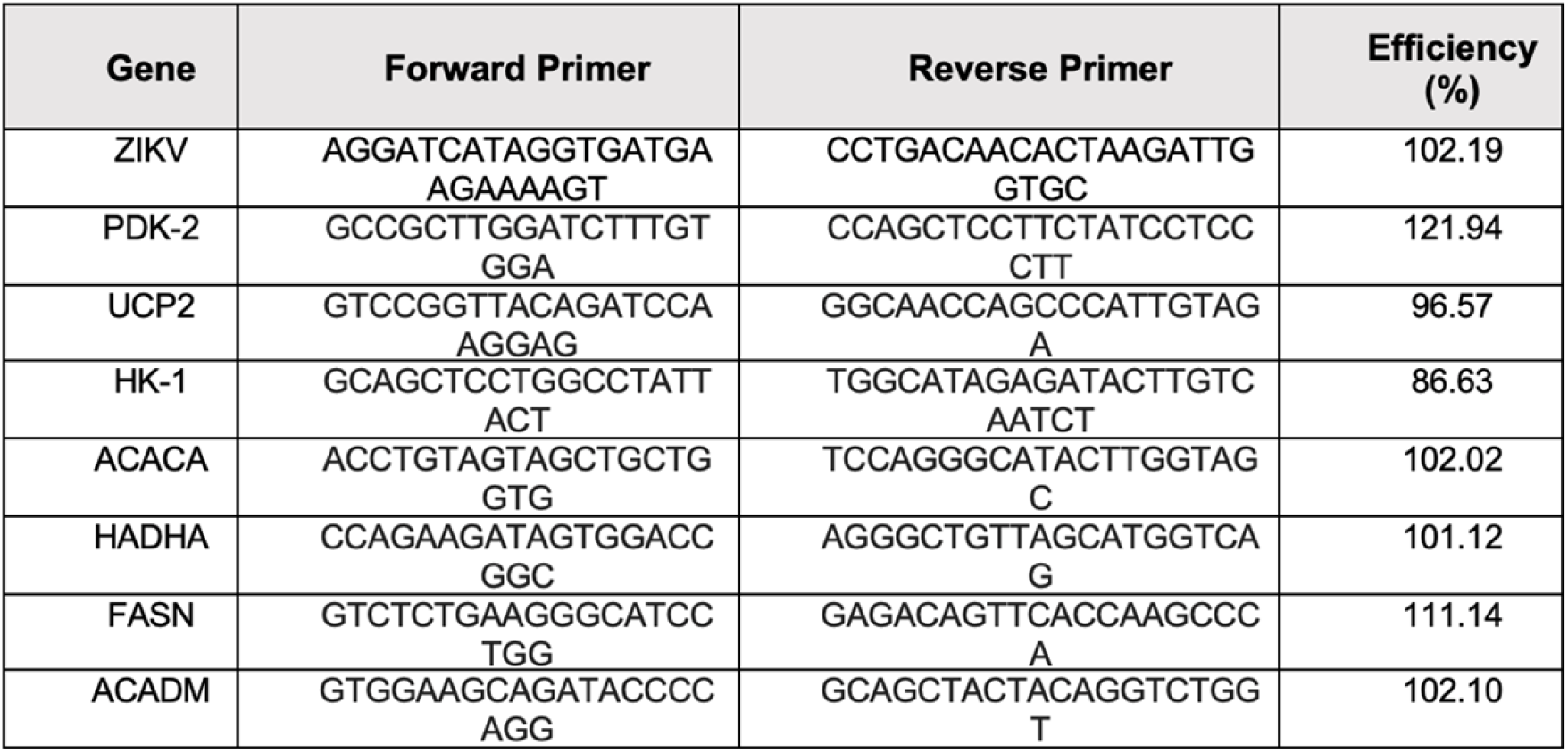
Efficiency and sequence of primers. Figure showing the sequence and calculated efficiency of the unpublished primers designed to assess the metabolic transcriptional profile of hi-NPC subtypes during exposure to ZIKV. Primers were concentrated at 100 nM and the annealing temperature at 64 °C.

